# The global fish and invertebrate abundance value of mangroves

**DOI:** 10.1101/2024.05.02.591889

**Authors:** Philine zu Ermgassen, Thomas A. Worthington, Jonathan R. Gair, Emma E. Garnett, Nibedita Mukherjee, Kate Longley-Wood, Ivan Nagelkerken, Kátya Abrantes, Octavio Aburto-Oropeza, Alejandro Acosta, Ana Rosa da Rocha Araujo, Ronald Baker, Adam Barnett, Christine M. Beitl, Rayna Benzeev, Justin Brookes, Gustavo A. Castellanos-Galindo, Ving Ching Chong, Rod M. Connolly, Marília Cunha-Lignon, Farid Dahdouh-Guebas, Karen Diele, Patrick G. Dwyer, Daniel A. Friess, Thomas Grove, M. Enamul Hoq, Chantal Huijbers, Neil Hutchinson, Andrew F. Johnson, Ross Johnson, Jon Knight, Uwe Krumme, Baraka Kuguru, Shing Yip Lee, Aaron Savio Lobo, Blandina R. Lugendo, Jan-Olaf Meynecke, Cosmas Nzaka Munga, Andrew D. Olds, Cara L. Parrett, Borja G. Reguero, Patrik Rönnbäck, Anna Safryghin, Marcus Sheaves, Matthew D. Taylor, Jocemar Tomasino Mendonça, Nathan J. Waltham, Matthias Wolff, Mark D. Spalding

## Abstract

Mangroves are a critical coastal habitat that provides a suite of ecosystem services and supports livelihoods. We undertake the first global analysis to estimate density and abundance of 37 commercially important fish and invertebrates that are known to extensively use mangroves. Geomorphic mangrove type, sea surface salinity and temperature, and length of mangrove forest edge were important in predicting the density of commercial fish and invertebrates, with deltaic systems supporting the highest densities. The model predicted high densities throughout parts of southeast Asia, the northern coast of South America, the Red Sea, and the Caribbean and Central America. Application of our model onto the global mangrove extent, estimates that mangroves support the annual abundance of nearly 800 billion young-of-year fish and invertebrates contained in our model. Our results confirm the critical role of mangroves globally in supporting fish and fisheries, and further builds the case for their conservation and restoration.

---

Food from the marine and coastal environment is a crucial source of protein and micronutrients for billions of people^1,2^. As the global human population and its requirement for food continues to increase, pressure for the marine and coastal environment to supply greater amounts of seafood will also increase^3,4^. In addition, marine fisheries support livelihoods for an estimated 260 million people, particularly in the Global South^5,6^. The production of marine and coastal food as well as the livelihoods supported are intrinsically linked to the condition of the environment, with healthy fish stocks dependent on the effective ecological functioning of freshwater, coastal and marine ecosystems^7^. Marine and coastal food production is therefore highly vulnerable to human driven environmental disturbance^7^, including over-harvesting, which has already driven losses in productivity and the collapse of some fisheries^8,9^. Coastal ecosystems have been highlighted as a critical driver of marine fisheries productivity^10^. In the face of widespread loss and degradation of these ecosystems, quantifying their role in marine fish and invertebrate production is key to help inform and support actions to sustainably manage these resources^11^.

Mangrove forests are complex and highly productive ecosystems, thriving in sheltered intertidal areas of tropical, subtropical and warm temperate coasts worldwide^12^. Their location, productivity and structure provide critical habitat for several finfish and shellfish species, giving shelter, food, solid structure for settlement, and critical nursery grounds^13,14^. For millennia they have supported human communities who fished and foraged in their waters, and today there are an estimated 4.1 million mangrove-associated fishers globally^15^. In addition to small-scale fisheries, mangrove-associated fisheries include some valuable export fisheries, notably of shrimp^16^ and crab species^17^. Across all fishery sectors, the importance of these fisheries is likely to grow in times of stress such as financial, social or economic instability, or particular environmental or climatic impacts. Well managed mangroves can support a relatively reliable and secure food supply enabling an important adaptive capacity for coastal communities. These fisheries benefits stand alongside other well-documented benefits such as carbon storage and sequestration^18^, protecting coastlines from storms and flooding^19^ and as habitat for birds, bats and other terrestrial species^20^.

Studies have focused on the role of mangrove seascapes in enhancing fisheries^21^ and the contribution of mangrove productivity to fish and shellfish biomass production^22^. Others have considered the role of mangroves in enhancing settlement and growth or hosting key life-history stages of individual species^23^. Despite these factors, there has been no attempt to derive a holistic global picture of fish/shellfish production associated with the world’s mangrove forests^24^. Here, we produce the first global analysis of the importance of mangroves to the density and abundance of commercially important fish and invertebrates that are known to utilise mangrove ecosystems extensively. This analysis gathered field data on the density of those species in mangrove areas, and used the Delphi technique^25^ to identify and weight expert knowledge on the biophysical drivers of the density of 37 mangrove-associated commercially exploited marine species, including species of fish, prawns, crabs and a bivalve. Geospatial data layers that describe the biophysical drivers were used to map the estimated density of each species for all locations where mangroves occur, and calculate the abundance associated with the world’s mangrove forests. To evaluate the importance of mangrove for subsistence and income for local livelihoods, the correspondence between our fish and invertebrate abundance estimates and the number of mangrove small-scale fishers is assessed.

## Drivers of mangrove fish and invertebrate density

A literature search and expert elicitation resulted in 481 field measurements of fish and invertebrate density (standardized to individuals 100 m^-2^), across 37 commercially important species (Supplementary Table 1). While many of these species have wide distributions, field data was geographically biased towards the Americas and to a lesser extent Asia, with limited data from southern and East Africa and the Middle East, and none for West Africa (Supplementary Fig. 1a). The subsequent models extend to the full range of all assessed species (often far beyond the location of the field data from the literature), and while the models show some species coverage for all mangrove areas, the distribution remains uneven, with important commercial species excluded, particularly from regions without primary data (Supplementary Fig. 1b - e). The greatest number of modelled species was centred on south and central America and the Caribbean, with up to 26 species represented in some areas. Data on bivalves were limited to species of mangrove cockle *Anadara tuberculosa*, which are confined to the Pacific coast of the Americas. In any one location there were up to two species of commercially important crabs, with richness centred in the Indian Ocean, while a maximum three species of penaeid prawns were distributed across the Indo-Pacific. However, no crustaceans could be modelled for the Atlantic-Eastern Pacific region.

**Table 1.** Top 10 countries for mangrove commercially-harvested marine species abundance. Different groups of species in billions per year, number in parentheses is the 95% confidence interval.

| Rank | All Species |  | Prawns |  | Finfish |  | Crabs |  | Bivalves |  |
| --- | --- | --- | --- | --- | --- | --- | --- | --- | --- | --- |
| 1 | Indonesia | 227<br>(41 - 1311) | Indonesia | 150<br>(25 - 915) | Indonesia | 76<br>(16 - 395) | Brazil | 36<br>(11 - 117) | Ecuador | 6<br>(2 - 20) |
| 2 | Malaysia | 74<br>(12 - 454) | Malaysia | 49<br>(8 - 310) | Brazil | 27<br>(7 - 99) | Madagascar | 21<br>(5 - 96) | Mexico | 4<br>(1 - 13) |
| 3 | Brazil | 63<br>(19 - 215) | Myanmar | 35<br>(6 - 219) | Malaysia | 25<br>(5 - 143) | Mozambique | 19<br>(4 - 92) | Colombia | 3<br>(1 - 11) |
| 4 | Myanmar | 53<br>(9 - 317) | Papua New Guinea | 34<br>(6 - 198) | Papua New Guinea | 19<br>(4 - 93) | Mexico | 14<br>(4 - 62) | Nicaragua | 2<br>(1 - 7) |
| 5 | Papua New Guinea | 53<br>(10 - 291) | Australia | 19<br>(3 - 108) | Myanmar | 18<br>(3 - 98) | Tanzania | 13<br>(3 - 59) | Honduras | 1<br>(0 - 4) |
| 6 | Madagascar | 39<br>(8 - 191) | India | 16<br>(2 - 106) | Mexico | 12<br>(4 - 39) | Venezuela | 8<br>(2 - 26) | Costa Rica | 1<br>(0 - 4) |
| 7 | Mozambique | 30<br>(6 - 148) | Viet Nam | 16<br>(3 - 93) | Australia | 10<br>(2 - 43) | Cuba | 5<br>(1 - 17) | El Salvador | 1<br>(0 - 3) |
| 8 | Mexico | 29<br>(9 - 113) | Madagascar | 12<br>(2 - 67) | India | 8<br>(2 - 45) | Colombia | 5 (1 - 25) | Panama | 1<br>(0 - 2) |
| 9 | Australia | 29<br>(6 - 151) | Pakistan | 10<br>(2 - 61) | Viet Nam | 7<br>(1 - 39) | Ecuador | 3 (0 - 23) | Guatemala | 1<br>(0 - 2) |
| 10 | Tanzania | 25<br>(5 - 123) | Tanzania | 8<br>(1 - 44) | Venezuela | 7<br>(2 - 27) | Kenya | 2 (0 - 7) | Peru | 0.03<br>(0 - 0) |

**Fig. 1.**
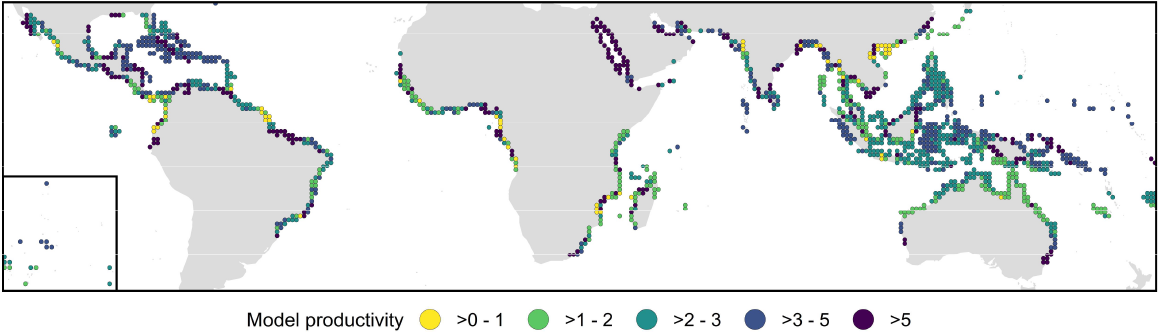
The density model output (individuals 100 m^-2^). Derived from the variables sea surface salinity (SSS), sea surface temperature (SST), mangrove edge (EDGE), and the correction factors for the gear type (G_j_) and the mangrove geomorphic type (C_k_). This output was visualised prior to applying individual species-specific density corrections and their presence/absence. Data summarised within 1° cells: inset shows central Pacific islands.

The mangrove fish and invertebrate density model contained the environmental variables sea surface salinity (SSS), sea surface temperature (SST), length of mangrove edge (EDGE). There were also correction factors for the species-specific density (A_i_), the gear type (G_j_) and the mangrove geomorphic type (C_k_) (Equation 1).

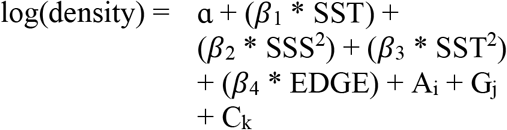

The base density model (i.e., prior to applying individual species corrections and their presence/absence) predicted high densities of fish and invertebrates with increasing sea surface salinity and in deltaic mangroves, with geomorphic setting having been identified as a key determinant of ecosystem function^26^. Mangrove edge length has been shown to be positively correlated with fish catches^21^; however, it was represented by a negative relationship in our model. On the map, the model predicted high densities of fish and invertebrates throughout parts of southeast Asia, particularly on the island of Borneo, large extents of the northern coast of South America, the Red Sea, and parts of the Caribbean and Central America (Fig. 1). Lower densities of fish and invertebrates were predicted for much of the coast of east Asia, central Africa and the Pacific coast of South America (Fig. 1).

## Regional patterns of mangrove fish and invertebrate density

The predicted densities of fish and invertebrates ranged from 0.09 to 11,019 individuals 100 m^-2^. These numbers were derived from the base density model, the species-specific modifiers, and the spatial variation in the numbers of modelled species. These maps illustrate broad patterns in the predicted density of fish and invertebrates in mangroves globally, and heterogeneity in density estimates both within and between regions. They are useful for examining density patterns within regions (see below), but are not appropriate for making large inter-regional comparisons because they are based on combined species totals and there is considerable spatial variation in the numbers of species modelled.

Along the Atlantic coasts of the Americas where our models have the greatest richness of commercially exploited finfish species (n = 26), densities at the national level were highest for some islands of the Caribbean such as Cuba (128 individuals 100 m^-2^, 95% CI: 50 - 360), as well as the Caribbean coast of Central America in Nicaragua (127 individuals 100 m^-2^, 95% CI: 44 - 425) and Belize (124 individuals 100 m^-2^, 95% CI: 47 - 357) (Fig. 2a). Geomorphic setting is considered a key factor structuring mangrove fish assemblages across the Americas (G.A. Castellanos-Galindo pers. comm.) and in this region hotspots of finfish density include the extensive mangrove coastline of Brazil to the east of the Amazon as well as the deltaic and lagoonal coasts of Colombia (Ciénaga Grande de Santa Marta) and Mexico, all with commercial finfish densities over 200 individuals 100 m^-2^ of mangroves per year (Fig. 2a). The model’s findings align with other observations that deltaic areas are characterised by high mangrove biomass and productivity^27^, and that within South America they represent critical areas of finfish abundance supporting local artisanal fisheries^28–30^.

**Fig. 2.**
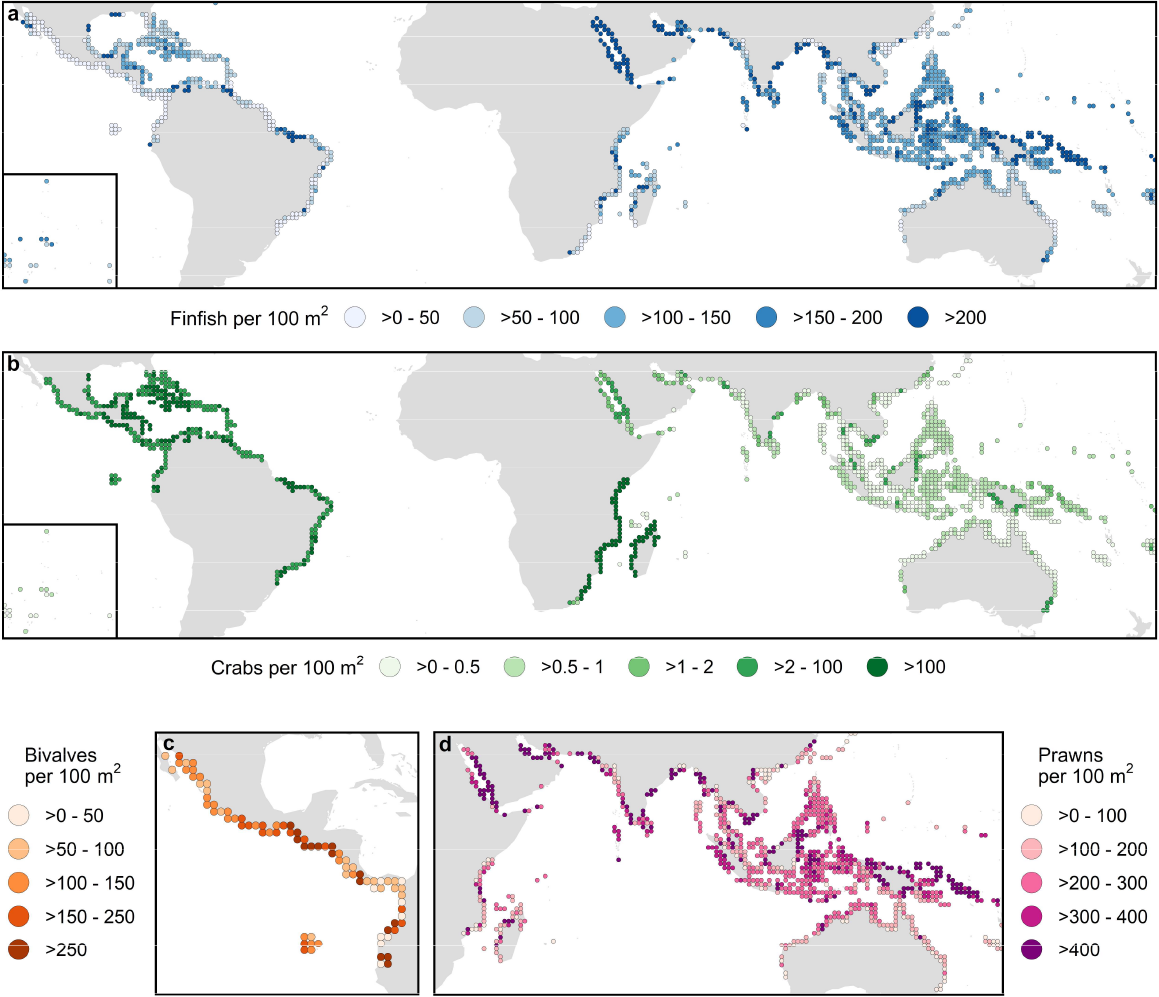
The modelled density of commercially important fish and invertebrate species due to the presence of mangrove ecosystems. Species grouped into (**a**) finfishes (n = 29), (**b**) crabs (n = 4), (**c**) bivalves (n = 1), and (**d**) prawns (n = 3). Although some species ranges also include West and Central Africa, predictions were removed for that region due to the lack of any field data to inform the modelling process. Data summarised within 1° cells: inset on (**a**) and (**b**) shows central Pacific islands.

Across the Indo-Pacific region there is a relatively even coverage of modelled finfish species and we see high predicted densities throughout the Red Sea and the Persian Gulf, as well as for countries such as Malaysia (418 individuals 100 m^-2^, 95% CI: 81 - 2274) and Papua New Guinea (339 individuals 100 m^-2^, 95% CI: 74 - 1627) (Fig. 2a). Within this region, finer scale patterns emerge with the deltaic coasts of northern Borneo, southern Viet Nam, and the south coasts of New Guinea among the areas supporting more than 200 finfish, and over 400 prawns, per 100 m^2^ of mangroves. Given the extensive areas of mangroves in many of these countries, the estimated overall abundance of commercial species is correspondingly high, and some of these mangroves have already been predicted to have a critical role in supporting large numbers of small-scale mangrove-associated fishers^15^. Elsewhere, areas which have already lost substantial areas of mangroves, such as across Java and much of the Philippines, have likely also lost a large proportion of these benefits^31^. The high finfish densities in this region are supported by the presence of *Atherinomorus lacunosus*, a small schooling species that had the highest predicted densities of any commercial finfish species in our model.

For crabs, density is high and quite consistent across the Atlantic and Pacific coasts of the Americas, due to the inclusion in our model of the two species from the genus *Ucides*, with the highest predicted densities in Brazil (283 individuals 100 m^-2^, 95% CI: 88 - 920), Venezuela (270 individuals 100 m^-2^, 95% CI: 79 - 983), Ecuador (190 individuals 100 m^-2^, 95% CI: 26 – 1,392) and Colombia (166 individuals 100 m^-2^, 95% CI: 38 - 845) (Fig. 2b). The two species, *U. cordatus* and *U. occidentalis*, are commercially and culturally important and a key part of the diet for coastal communities in the region^32,33^, with densities driven by factors such as tidal influence, mangrove condition and fishing pressure operating at a range of spatial scales^34,35^. By contrast, across the Indo-Pacific, crab density (species *Scylla serrata* and *Neosarmatium africanum*) appears to be highly heterogeneous and lower than in the Americas. High densities in the Western Indian Ocean, including Tanzania (919 individuals 100m^-2^, 95% CI: 206 - 4158) Madagascar (729 individuals 100m^-2^, 95% CI: 164 - 3267), Mozambique (668 individuals 100m^-2^, 95% CI: 145 - 3114), and Kenya (334 individuals 100m^-2^, 95% CI: 83 - 1365), are driven by the inclusion of an additional species in the model, *N. africanum*, that is restricted to this region (Supplementary Fig. 1c).

Three species of penaeid prawns were included, with ranges covering much of the Indo-Pacific (Supplementary Fig. 1d). Several studies have highlighted a correlation between the area or linear extent of mangrove habitat, as well as factors such as sea surface temperature and latitude, and the commercial catches of penaeid prawns, with mangroves acting as a nursery habitat for post-larval prawns^16,36,37^. Within the model the penaeid prawns were predicted to have some of the highest densities of any species. This high abundance coupled with a high market value means that penaeid prawns are considered an economically important mangrove-associated fisheries commodity^36^. High densities of prawns are modelled for the Middle East and Arabian Sea regions (for example Pakistan (1,298 individuals 100m^-2^, 95% CI: 219 – 7,735) and Saudi Arabia (1,110 individuals 100m^-2^, 95% CI: 140 – 8,946)) (Fig. 2d), driven by the presence of *Penaeus merguiensis* and *P. indicus*. These two species are highly dependent on mangrove ecosystems to complete their life history, being observed almost exclusively in mangrove creeks^16^. Despite these high densities, the relatively low overall extent of mangroves in this region means that the highest overall abundance of prawns are represented elsewhere (Table 1).

On the Pacific coasts of the Americas the model includes few finfish and no prawns. However, this region comprises the only bivalve species *Anadara tuberculosa* in our model. *A. tuberculosa* is distributed across ten countries of the Pacific seaboard of the Americas and showed the highest densities in Ecuador (345 individuals 100 m^-2^, 95% CI: 101 – 1,188) and Costa Rica (252 individuals 100 m^-2^, 95% CI: 78 - 817) (Fig. 2c), where they represented nearly 60% of the total density of individuals. *A. tuberculosa* is associated with the root system of red mangrove (*Rhizophora mangle*) and represents one of the commercially most important species in the region^38^. Other commercially important bivalve species, e.g., *A. similis, Crassostrea* spp. and *Geloina* spp. are widely harvested from mangroves around the world, but could not be incorporated into our model due to a lack of species specific density data.

### Fish and invertebrate abundance from mangroves

Application of our model onto the 2020 global mangrove extent^39^, estimates that the presence of mangroves supports an annual abundance of nearly 800 billion (95% confidence intervals (CI): 160 – 4,200 billion) young-of-year fishes, prawns and bivalves, and adult crabs from across the commercially important species considered here. It should be noted that abundance at the early life-history stage, and across a range of species with very different life-history parameters does not directly equate to potential economic or biomass gains. Furthermore, such numbers represent a substantial under-estimate of the commercial importance of mangroves across all commercial species, focusing only on a subset of species targeted by fisheries, and with no data at all for abundance from West and Central African mangroves. Of the global 800 billion total, the three species of the genus *Penaeus* represented nearly half (47.9%) the total, with the 29 species of finfish contributing a further 32.6%. The remaining amount was split between four species of crabs (17.1%) and the single bivalve, *A. tuberculosa*, (2.4%).

The contribution of different groups to the total abundance world-wide is affected by species data availability. For example, prawn species are an important component of both mangrove ecosystems and fisheries catches across the Americas^40,41^, and their inclusion would greatly increase numbers in this region; however, penaeids quite strongly influence the model numbers across much of the Indo-Pacific where available data met the requirements for inclusion in the model. By contrast, crabs make up a larger proportion of total numbers in the Americas and in east and southern Africa (Fig. 2b), while included bivalves are highly restricted to the Pacific coast of the Americas. Given this regional variation in data availability, the description and exploration of the model and its predicted patterns focuses at regional (or finer) scales where there are similar taxa and comparable species richness within the model.

Southeast Asia supports some of the highest abundance of fish and invertebrates, with over half of the global numbers predicted by our model, which includes 51.5% of the global finfish and nearly 70% of the global prawn numbers. Three species of prawns account for two thirds of the total numbers in Southeast Asia, but fish are also a large component with *Atherinomorus lacunosus*, and *Gerres filamentosus* being among the most numerous. In addition to these highly abundant species, the model also contains species such as the mangrove red snapper (*Lutjanus argentimaculatus*) that, although making up only a small proportion of the total numbers, represents a particularly high-value species important across the region^42^. Given its extensive mangrove habitat, Indonesia has the highest totals, with this abundance shown to support livelihoods for communities across the country^43,44^ and to have a critical role in food security and nutrition^45^. Similarly high abundance of commercially important mangrove associated fish and invertebrate occurs in Malaysia, Myanmar and Papua New Guinea (Table 1).

Among the modelled species, crab abundance is high across the mangrove coasts of the Americas, underpinned by the high-densities of *U. cordatus* and *U. occidentalis*. Multiple studies have highlighted the socio-economic importance of crab and bivalve fisheries for local use across Central and South America^35,46,47^. In the western Indian Ocean, total abundance is dominated by two crab species (*S. serrata* and *N. africanum*) (56.4%) This includes one crab species, *N. africanum*, that is restricted to this region and has the highest modelled density of any species in our models. Crabs are highly important species here, not only for local consumption. In Madagascar, small-scale subsistence *S. serrata* fisheries have shifted to market driven exports, resulting in a fivefold increase in their price^48^. In addition, crabs and other invertebrate species may fulfil important ecosystem engineering functions due to their low functional redundancy^49^.

In contrast to other regions, total abundance of commercially important mangrove associated fish and invertebrates for large parts of the Middle East remains low. Although the models predict high density per unit area, mangroves in much of this region are limited in total extent. Moving across South Asia, the pattern changes, with highly heterogeneous numbers, including very high numbers in low-lying coastal areas around rivers with extensive mangroves. Data gaps for many key species reduce regional and global totals, and constrain use in broad regional comparisons, however, the patterns that our model highlights are important for a better understanding the value of mangrove ecosystems. Indeed, many of the species we were unable to model or map likely follow similar patterns to those modelled.

### Mangroves’ contribution to livelihoods and global food security

Mangrove fisheries provide important contributions to the provision of dietary protein^50^ as well as subsistence, recreation, and employment in commercial fisheries^15,51^, known to involve numerous stakeholders between fishers and markets^52^. The correspondence between our abundance estimates and the number of fishers participating in in-mangrove, near-shore subsistence and artisanal and near-shore commercial fisheries^15^ (hereafter referred to as small-scale fishers) showed considerable heterogeneity, but with some distinct regional variation (Fig. 3a). Areas predicted to support high fish and invertebrate abundance, and large numbers of small-scale fishers^15^ were centered on

**Fig. 3.**
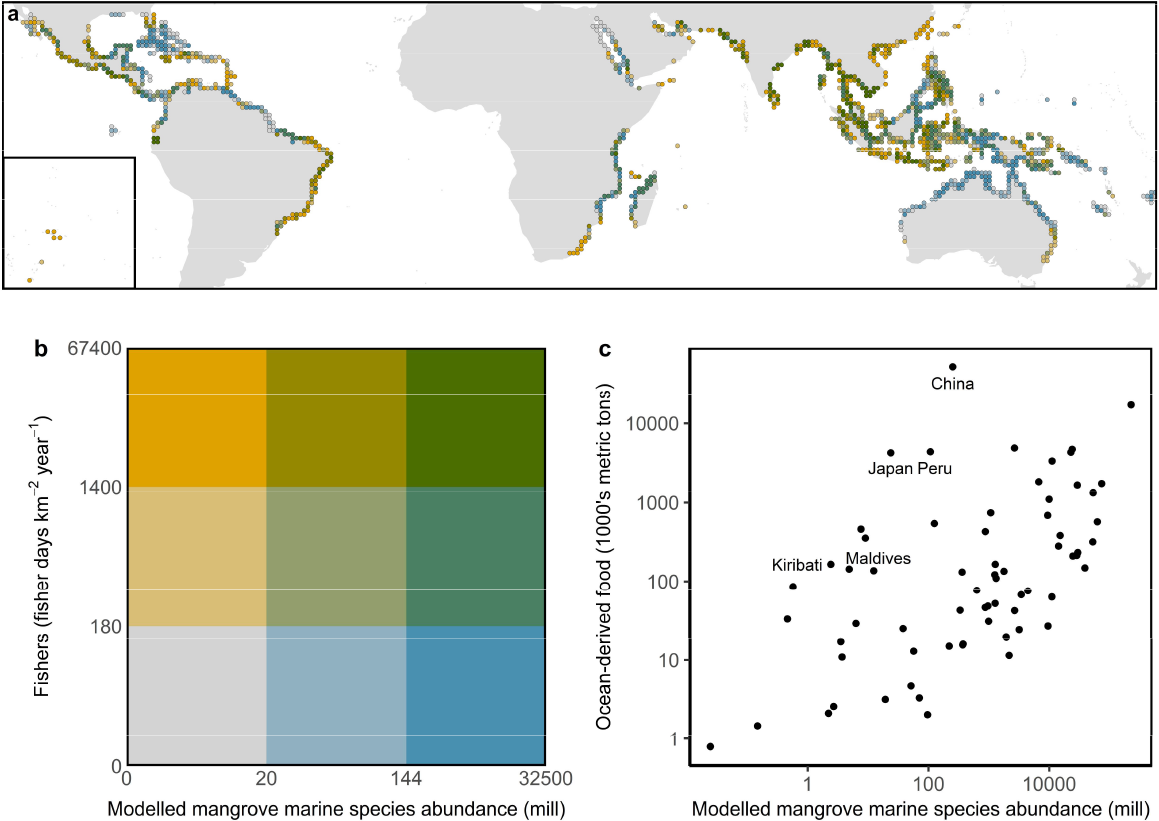
Mangroves’ contribution to livelihoods and global food security. (**a**) The correspondence between modelled mangrove commercially-harvested marine species abundance and the number of fishers participating in in-mangrove, near-shore subsistence and artisanal and near-shore commercial fisheries^15^. Data summarised within 1° cells, inset shows central Pacific islands, (**b**) key to colours in panel (a) showing the breakpoints in terms of modelled mangrove fish and invertebrate abundance and fishers for the different categories, and (**c**) the relationship between a country’s mangrove commercial fish and invertebrate abundance (this model) and its total ocean-derived food production. Countries highlighted in the text are labelled.

South and Southeast Asia, particularly in highly populated areas such as the Mekong delta. Conversely, relatively low values of those two variables were apparent in the Caribbean, northern Red Sea and parts of Australia. Areas that had relatively low fish and invertebrate abundance but still supported high numbers of small-scale fishers were evenly distributed across the world (Fig. 3a). By contrast, areas of high abundance and low numbers of small-scale fishers can be seen across much of tropical Australia, Southern Papua (Indonesia and Papua New Guinea) as well as the Western Caribbean (Cuba, Bahamas, South Florida), although it should be noted that in certain regions such as the Western Caribbean and Northern Australia this abundance supports high value recreational fisheries^51^.

The national fish and invertebrate abundance from mangrove ecosystems predicted by our model was highly positively correlated with a country’s total ocean-derived food production^53^ (*t*_*65*_ = 5.2, *P* < 0.001, r^2^ = 0.546; Fig. 3c). Outliers within this trend were Peru, Japan and China, as well as smaller island nations such as Kiribati and the Maldives. These countries have high total ocean-derived food production, but only small mangrove areas and thus relatively less mangrove associated fish and invertebrate abundance. To date, the relationship between mangrove ecosystems and enhanced coastal and inshore fisheries has largely been assumed, with empirical relationships between mangrove descriptors e.g., area or edge length, and fish catches based on certain species or countries^22^. The overall pattern found in this study supports the inference that mangroves play a relatively consistent and important role in global fisheries and as such adds another supporting argument for their conservation and restoration, alongside other quantified benefits such as carbon storage or coastal flood protection.

## Conclusions

Mangrove forests are known to provide important nursery areas for finfish and invertebrate species of commercial importance^13,14^. While a number of studies have attempted to quantify the enhancement role of mangroves for commercially important species, there is insufficient data to measure such enhancement at the global scale for all commercial species. The current work thus focuses on a subset of commercially important fish and invertebrate species for which data are available and which are known to have a high dependency on mangroves, and then models the likely density within the mangroves. The model outputs confirm the high densities of commercially important fish and invertebrate species in many mangrove areas world-wide. Within particular regions where the model includes a consistent set of modelled species, the generated maps enable within-region comparison. Our study highlights the particular role of deltaic coasts in Southeast Asia and the Americas, and also arid coastlines of the Middle East in supporting high densities of commercial species per unit area. Clearly, overall abundance of commercially important fish and invertebrate species is strongly influenced by overall mangrove extent, and in this regard, the mangroves of Indonesia support the greatest abundance of commercial fish and invertebrates of any single country in the Indo-Pacific, with Brazil and Mexico dominating total abundance in the Americas. Particular attention can also be drawn to important ecological and geomorphological features which are promoting high numbers of commercial species, and which will clearly benefit from coherent management such as the deltaic coastlines of the Sundarbans, southern New Guinea, north-east Borneo and the northern coasts of South America.

The importance of commercial mangrove fish and invertebrates to small-scale and local fishers has been emphasised in other work^15^, but overlapping such information with models of abundance highlights areas where fishing pressure and fish abundance may be interacting, with areas of low commercial fish abundance and high apparent fishing effort being areas in need of particular attention from a sustainability and fisheries management perspective. A close correlation between the estimated abundance of fish supported by mangroves from our models and total fisheries dependence at the national level highlights the critical importance of mangroves to fisheries and food security world-wide.

## Methods

### Fish and invertebrate density data

A literature search was undertaken on 20th September 2016 in Scopus using the following search terms to search for relevant titles, abstracts or keywords ((mangrove*) AND (densit*) AND (*fish* OR invertebrate* OR crab* OR mollusc* OR shrimp* OR mussel*)) to identify papers containing species-specific fish and invertebrate data from field studies undertaken in mangroves (n = 520). The search was repeated on 3rd June 2021 to capture more recent papers reporting relevant values. Additionally, participating experts were asked to contribute published datasets, and relevant references were searched for mention of other possibly relevant publications in a snowball approach.

The abstracts of all identified papers were screened to assess their relevance, and relevant papers were then searched for extractable fish and invertebrate density data. Papers providing data from areas outside of the native range of mangroves (e.g., Hawaii), were excluded. All commercial fish, bivalve and crustaceans data were extracted, with values standardised to represent the number of individuals 100 m^-2^. Only gear types that provided information as to the spatial area of sampling were included. Gear type and standard error were also extracted into the dataset. Where standard deviation was provided, this was transformed into standard error. Gear types included trawl, quadrat, mark-recapture, visual survey, block net, seine net, and lift net. Mark-recapture data were included from a single study^54^ in which the authors converted the data to abundance via the Chao model. To avoid pseudoreplication within the dataset all samples from the same estuary, or fewer than 5 km apart, were aggregated to a single mean value. Similarly, where papers had recorded data at multiple timepoints, annual averages were derived when possible.

### Spatial data

#### Spatial extent

A grid with a spatial resolution of 1 km^2^ was created for the entire world. Grid cells (n = 488,097) that contained mangroves between the years 1996 and 2020 from the Global Mangrove Watch (GMW; v3.14) time series^39^ were identified, and formed the spatial framework of the analysis.

#### Environmental covariates

To estimate the fish and invertebrate density within mangroves, a statistical model that linked the fish and invertebrate density data to environmental covariates was created. The environmental covariates were identified through a Delphi approach conducted over a two-month time frame (May 19 - July 20, 2016). Seventeen mangrove fish and/or invertebrate experts from across the globe, with a mean 17 years experience with mangroves and field experience in 39 countries and territories were asked to determine which abiotic and human factors most influenced the density of mangrove-affiliated fish and invertebrates. The countries in which of experts had experience in included Australia (n = 7), Papua New Guinea (4), Singapore (4), Fiji (3), Malaysia (3), New Caledonia (3), Tanzania (3) and Thailand (3). Fourteen of the experts identified themselves as having expertise in fish and/or invertebrate ecology, with the remaining three identifying their expertise as fisheries (catch), food-webs and spatial ecology. The experts were asked to provide insight on three aspects of mangrove fisheries: (1) parameters that might influence fish and invertebrate density, (2) the direction of the relationship between the parameter and fish and invertebrate density, and (3) the geographic scale at which the parameter might exert the strongest influence. Experts ranked the options on a Likert scale of 1–5^55^. Factors which reached consensus as being important in determining fish and invertebrate density in mangrove systems were taken forward and considered for inclusion in the global model of mangrove-affiliated fish and invertebrate densities. Consensus was reached when more than 75% of experts scored the factors as 4 or 5 on the Likert scale.

Following two rounds of the Delphi technique, consensus was reached on ten of the initial 45 proposed factors (Supplementary Table 2): mangrove extent, mangrove geomorphic type, duration of inundation, fish and invertebrate species mangrove-dependency, mangrove condition, proximity to seagrass (where cooccurring), fishing pressure, mangrove edge length, extent of estuary/embayment proximate to the mangrove, and fish and invertebrate species probability of occurrence. Global geospatial datasets representing these factors were sought (Table 2); however, suitable datasets representing duration of inundation or proximity to seagrass were not available on a global scale. While consensus was not reached on the importance of sea surface temperature, salinity, and net primary productivity to fish and invertebrate density within mangroves specifically, they are known to influence fish abundance at a large spatial scale^56,57^, and were therefore retained as covariates for the model fitting.

**Table 2.** The environmental covariates included the global model of mangrove-affiliated fish and invertebrate densities.

| Variable | Rationale | Data Resolution | Model Fitting Data | Model Prediction Data | Ref |
| --- | --- | --- | --- | --- | --- |
| Geomorphic Type | Coastal geomorphology is a key factor that structures the relationship between habitat and coastal fish communities<sup>63</sup>. For example, coastal mangroves and estuarine mangroves support a different suite and abundance of tropical prawn species<sup>64</sup>. | NA | 1996, 2007-2010 and 2015 - 2020 | 1996 to 2020 Composite | <sup>65</sup> |
| Mangrove Condition | Mangrove forest habitat degradation can result in lower fish catch rates<sup>31</sup>. | NA | Static | Static | <sup>39</sup> |
| Mangrove Area | Areal extent of mangroves is known to positively impact the abundance of mangrove associated fish and invertebrate species across a range of geographies<sup>66-68</sup>. | 25 m | 1996, 2007-2010 and 2015 - 2020 | 2020 | <sup>39</sup> |
| Edge Length | Mangrove edge habitats are often found to harbour higher densities of fish than sites within mangroves and therefore likely play a disproportionately important role in total fish abundance<sup>69,70</sup>. | 25 m | 1996, 2007-2010 and 2015 - 2020 | 2020 | <sup>39</sup> |
| Tidal Amplitude | The amount of intertidal habitat available, which is a function in part of the tidal amplitude, is an important factor in explaining tropical prawn abundance<sup>71</sup>. | ~7 km<sup>2</sup> | Static | Static | <sup>72</sup> |
| Coastal Embayment Area | The size of the estuary associated with mangroves has been found to influence the abundance of fish and prawns. | NA | Static | Static | <sup>65</sup> |
| Fishing Pressure | Fish species richness and abundance is lower in areas with high fishing pressure<sup>73</sup>. | 1 km<sup>2</sup> | Static | Static | <sup>74</sup> |
| Proximity to Seagrass | Fish have been shown to utilise mangrove habitats near seagrass beds when the tide allows. There is therefore a positive synergistic effect between seagrasses and mangroves across the tidal cycle where they co-occur<sup>75-77</sup>. | No data |  |  |  |
| Duration of Inundation | Mangrove areas that are underwater for longer periods provide habitat to fish and aquatic invertebrate species for a greater proportion of the day<sup>78</sup>. | No data |  |  |  |
| Sea Surface Salinity | Sea surface salinity is often used in modelling large scale fish abundance<sup>79,80</sup>. | 25 km<sup>2</sup> | 2011 - 2020 | 2020 | 81 |
| Sea Surface Temperature | Sea surface temperature is commonly considered in large scale fisheries modelling as an important variable explaining fish abundance<sup>82,83</sup>. | ~5.55 km<sup>2</sup> | 2001 - 2020 | 2020 | 84 |
| Net Primary Productivity | Primary productivity explains variability in fish production in global and tropical seas<sup>82,85</sup>. | ~18 km<sup>2</sup> | 2002 - 2002 | 2020 | 86 |

The environmental covariates were represented by two types, firstly static variables whose value did not change over the period of the fisheries data, and time series variables whose value could be more closely attributed to an individual fisheries survey year (Table 2). For static variables the single value was used both for model fitting and predictions. For the time series variables, the value used for model fitting was from the year closest in time to the collection year of each fish survey data point, with the model predictions using the 2020 data to match the most recent global mangrove extent. For raster datasets, to determine the environmental covariate value for each grid cell, the centroid was used. The raster value closest to the grid cell centroid for the variables sea surface salinity (SSS), sea surface temperature (SST), tidal amplitude and net primary productivity (NPP) was identified. The spatial data processing was carried out in ArcGIS, and R^58^ using the packages sp^59^, raster^60^, ncdf4^61^ and rgdal^62^.

##### Geomorphic Type and Mangrove Condition

The geomorphic type of the mangroves in each 1 km^2^ cell was derived from a mangrove biophysical typology^65^. This typology was developed for a different mangrove extent, therefore it was updated to match the spatial extent of GMW v3.14^87^. The typology assigns areas of mangroves into ‘units’ based on their proximity to macroscale coastal features which determines their geomorphic class – deltaic, estuarine, lagoonal and open coast. For cells that intersect multiple mangrove typological units, the geomorphic type of the unit covering the largest proportion in a cell was used. The condition of mangroves in a cell was assessed at the scale of the typological units. The dominant (i.e., the unit covering the largest proportion of a cell) typological unit within a cell was identified, with condition the percentage change in mangrove area of that unit between 1996 and 2020. For those units with an infinite percentage increase in area i.e., those with area = 0 in 1996 and area > 0 in 2020, the value was set to the maximum percentage from the other units.

##### Mangrove Area

The area of mangrove within each of the 488,097 cells was calculated for all years (1996, 2007-2010 and 2015 – 2020) within the GMW v3.14 dataset^39^. Given the misregistration identified in the Japan Aerospace Exploration Agency Synthetic Aperture Radar mosaics that result in inflated change statistics for the GMW dataset, we calculated the adjusted area based on equation three and the commission and omission values from Table 10 in Bunting et al.^39^. Model predictions were mapped onto the cells that intersect with the 2020 extent (n = 477,199).

##### Mangrove Edge Length

The edge length was derived by converting the mangrove extent polygons (1996, 2007-2010 and 2015 – 2020) of the GMW v3.14 dataset^39^ to polylines. The length of mangrove edge (in meters) within 2km a buffer of each 1km grid cell was calculated. For estuarine deltaic, and lagoonal mangroves, the length of all edges within 2km was used; however, for open coast mangroves, this approach included a disproportionate amount of landward mangrove edge that would have no mangrove fisheries density value. Therefore, for open coast mangroves, only the mangrove edge lengths that fell within a 100m landward buffer from the shoreline (derived from Database of Global Administrative Areas v3.6, https://gadm.org/index.html) were used. While the mangrove extent data are available at moderate resolutions (25 m), it likely underestimates the extent of small watercourses within the mangroves. Model predictions used the 2020 mangrove extent to quantify edge length.

##### Sea Surface Salinity

Data from the European Space Agency’s Sea Surface Salinity Climate Change Initiative^81^ were accessed. Monthly mean data centred on the 1^st^ and 15^th^ day of each month covering the full assessment period, January 1^st^ 2011 to September 15^th^ 2020, were used. The 234 netCDF-4 were imported into R, and the coordinate system was converted from the Equal Area Scalable Earth grid to rasters with a World Geodetic System 1984 coordinate system. Ten mean annual SSS composites were created and linked temporally to the fish density data, with the predictions mapped onto the 2020 composite.

##### Sea Surface Temperature

Daily average ocean surface temperature adjusted to a standard depth of 20 cm were accessed from the Copernicus Climate Change Service^84^. Data were Level 4 spatially complete global sea surface temperature based on measurement from multiple sensors. Daily data in netCDF-4 form for the 1^st^ and 15^th^ of each month between 2001 and 2020 (n = 480) were downloaded and imported into R. Twenty mean annual average SST composites were created, with the predictions mapped onto the 2020 composite.

##### Net Primary Productivity

Net primary productivity data based on the Vertically Generalized Production Model (VGPM)^86^ were downloaded. The VGPM model provides an estimate of NPP, based on chlorophyll from MODIS data, available light and photosynthetic efficiency. Two hundred and twenty monthly files covering 2002 to 2020 in .xyz format were imported into R and converted to rasters. Nineteen mean annual NPP composites were created, with the predictions mapped onto the 2020 composite.

##### Tidal Amplitude

Tidal amplitude data were based on the Finite Element Solution tide model, FES2014. FES2014 integrates altimeter data from multiple satellites into a 2/3-D ocean hydrodynamics model^88^. A previous iteration (FES2012) was assessed as being one of the most accurate tide models for shallow coastal areas^88^, with significant improvements in predictions for those areas identified in FES2014^72^. For the analysis the principal lunar semi-diurnal or M2 tidal amplitude was used, as in most locations this is the most dominant tidal constituent^89^. The tidal amplitude raster with a pixel resolution of 1/16° (∼7km^2^ at the equator) was downloaded. This static data layer was used for both model fitting and model predictions.

##### Coastal Embayment Area

Coastal embayment area was assessed based on spatial polygons representing geomorphic features along the coastline. These spatial polygons were the framework for the mangrove typology (see Worthington et al.^65^ for full details) and identified based on rapid change in direction of a high-resolution coastline. These polygons represent individual riverine estuaries and deltas, or coastal lagoons and bays. The area of coastal embayment polygons within a 2km buffer of each grid cell was calculated.

##### Fishing Pressure

Spatial data on fishing effort (boat-meters per km^2^) within the coastal zone was used. Fishing effort data (number of boats, the length of boats, and the spatial boundary of the fishery) was extracted from FAO country profiles, published and grey literature and distributed across the coastal zone using contextual information on the distance from shore, distance from port, and the depth of the fishery^74^. Six regional raster datasets (pixel resolution 1km^2^) were combined and the raster value closest to the grid cell centroid was calculated. Owing to missing data in certain regions, distances between the grid cells and the fishing effort layer could be extremely large. Therefore, any grid cell > 100 km from the fishing effort dataset had its value set to 0.

### Data analysis

A linear model using generalized least squares from the package nlme^90^ was fitted to the fish and invertebrate density data in R^58^. The density data was log transformed to reduce the impact of extreme values. An initial model was fitted including the following variables: sea surface salinity, sea surface temperature, net primary productivity, geomorphic type, mangrove condition, mangrove area, edge length, tidal amplitude, coastal embayment area and fishing pressure. In addition, to reduce extreme density predictions at high values of sea surface salinity and sea surface temperature, the square of these variables was included.

Examination of the residuals of the initial model suggested that it violated the assumption of homogeneity of variance, and therefore the inclusion of different variance structures was tested^91^. A structure that allows different variances per stratum of the variable geomorphic type produced the lowest AIC; however, there was also an indication of a difference in variance across values of mangrove edge. Therefore, a combined structure that allows different variances per stratum of geomorphic type and power of the variance covariate for mangrove edge was used.

The model containing all variables and the variance structure was refined over multiple iterations, by dropping the least significant variable at each step. Gear type and species were included as factor variables, and were retained throughout the model fitting process. In the case of gear type, all pairwise gear comparisons were considered and gear types not significantly different from one another were merged. The mangrove geomorphic type was retained in the final model, along with the variables SST, SST^2^, SSS^2^, and edge length.

The model was then predicted over the spatial data framework for the cells within the grid which contained mangrove in 2020 (n = 477,199), using the cell values for geomorphic type, SST, SSS, and edge length. For the predictions the correction factor for gear type was set to 1. For certain areas the combination of covariates resulted in density predictions for this predictive surface greatly above those based on the covariate values represented within the input fish and invertebrate density field data. Therefore, any cells with predictions greater than the maximum prediction of the fisheries model were removed (n = 6,973). In addition, a further 285 cells from Hawaii and French Polynesia where mangroves have been introduced were removed. Ninety-five percent confidence intervals were created for all predictions using 1.96 * standard error of the model fit.

The model output was predicted densities (individuals 100m^-2^) for each of the 37 species across the 470,226 grid cells. To assure that the model did not predict species’ densities outside their native ranges we used Aquamaps^92^ to constrain species predictions. The current distribution for the 37 species were downloaded and converted to presence/absence maps using a threshold 0.01, representing all areas in which any given species was predicted to be present. These presence/absence maps were then multiplied by the density values to remove density predictions outside the species’ native range. Aquamaps was used for all species apart from the recently described *N. africanum*. For *N. africanum* the range was set as the east coast of Africa from Natal in South Africa to the middle of Somalia, and the whole of Madagascar^93^. Owing to the lack of fish and invertebrate density data from West and Central Africa (Supplementary Fig. 1a), this region was removed from this analysis.

To examine differences between types of fish and invertebrates, the 37 species were grouped into fish, crab, bivalves and prawns. To calculate the abundance of the 37 species in mangrove areas, the cell species densities (individuals 100 m^-2^) were multiplied by the area of mangroves within each cell in 2020. To evaluate mangrove contribution to employment, the correspondence between the fish and invertebrate abundance value of mangroves, and the number of fishers participating in in-mangrove, near-shore subsistence and artisanal and near-shore commercial fisheries (small-scale fishers) was assessed. As such a separate data source on fishing pressure than that included in the fish and invertebrate model (see above) was used. Data on the intensity of small-scale fishing (fisher days km^-2^ year^-1^) from zu Ermgassen et al.^15^ was spatially matched to our grid cells (n = 392,634). Data was summarised to 1° cells and the R package biscale^94^ was used to split the data into nine groups representing combinations of low, moderate and high values of total fish and invertebrate abundance, and small-scale fishers. To assess the contribution of mangroves to overall ocean-sourced food, we correlated our estimates of the national fish and invertebrate abundance value of mangroves with data on total ocean derived food production using data from the Food

Balance Sheets produced by the Food and Agriculture Organization of the United Nations^53^. Data were available for 67 countries.

## Data availability

The raw fish density data, and species and species group predictions are available on Zenodo (https://doi.org/10.5281/zenodo.11097214).

## Code availability

Code used in this study is available on Zenodo (https://doi.org/10.5281/zenodo.11097214).

## Acknowledgements

This work forms part of a project supported by the International Climate Initiative (IKI). The German Federal Ministry for the Environment, Nature Conservation and Nuclear Safety (BMU) supports this initiative on the basis of a decision adopted by the German Bundestag. Initial work on this study was supported by the Lyda Hill Foundation. The authors would like to thank Hederick Dankwa, Neil Loneragan, Roland Nathan Mandal, and Lawrence Rozas for input at an early stage in the project.

## Author Contributions

PE, TAW, and MDS conceived the study. NM, KA, OAO, RB, AB, RBe, JB, GACG, VCC, RMC, DAF, CH, NH, RJ, JK, UK, SYL, BRL, CNM, ADO, CLP, AS, MS, MDT and NJW collected the data underpinning the analysis. PE, TAW, JRG, NM, KLW developed the methodology, with TAW and JRG and conducting the statistical modelling. PE, TAW, EEG, KLW, TG and IN curated the data. PE, TAW, IN, RB, and MDS wrote the first version of the paper, with all authors providing input into subsequent versions of the manuscript.

## Competing interests

The authors declare no competing interests.

## Notes

DOI:10.5281/zenodo.11097214

### Competing Interest Statement

The authors have declared no competing interest.

## References

1. Golden, C. D. et al. Nutrition: Fall in fish catch threatens human health. Nature 534, 317–320 (2016).

2. FAO. The State of World Fisheries and Aquaculture 2020. Sustainability in action. (FAO, 2020). doi:10.4060/ca9229en.

3. Free, C. M. et al. Expanding ocean food production under climate change. Nature 605, 490–496 (2022).

4. Costello, C. et al. The future of food from the sea. Nature 588, 95–100 (2020).

5. Short, R. E. et al. Harnessing the diversity of small-scale actors is key to the future of aquatic food systems. Nat. Food 2, 733–741 (2021).

6. Teh, L. C. L. & Sumaila, U. R. Contribution of marine fisheries to worldwide employment. Fish Fish. 14, 77–88 (2013).

7. Cao, L. et al. Vulnerability of blue foods to human-induced environmental change. Nat. Sustain. 6, 1186–1198 (2023).

8. FAO. The State of World Fisheries and Aquaculture 2020. Towards Blue Transformation. (Food and Agriculture Organization of the United Nations, 2022).

9. Palomares, M. L. D. et al. Fishery biomass trends of exploited fish populations in marine ecoregions, climatic zones and ocean basins. Estuar. Coast. Shelf Sci. 243, 106896 (2020).

10. Barbier, E. B. et al. The value of estuarine and coastal ecosystem services. Ecol. Monogr. 81, 169–193 (2011).

11. Nordlund, L. M., Unsworth, R. K. F., Gullström, M. & Cullen-Unsworth, L. C. Global significance of seagrass fishery activity. Fish Fish. 19, 399–412 (2018).

12. Spalding, M. D., Kainumu, M. & Collins, L. World Atlas of Mangroves. (Earthscan, 2010).

13. Jänes, H. et al. Quantifying fisheries enhancement from coastal vegetated ecosystems. Ecosyst. Serv. 43, 101105 (2020).

14. Anneboina, L. R. & Kavi Kumar, K. S. Economic analysis of mangrove and marine fishery linkages in India. Ecosyst. Serv. 24, 114–123 (2017).

15. zu Ermgassen, P. S. E. et al. Fishers who rely on mangroves: Modelling and mapping the global intensity of mangrove-associated fisheries. Estuar. Coast. Shelf Sci. 248, 106975 (2020).

16. Manson, F. J., Loneragan, N. R., Skilleter, G. A. & Phinn, S. R. An evaluation of the evidence for linkages between mangroves and fisheries: A synthesis of the literature and identification of research directions. Oceanogr. Mar. Biol. An Annu. Rev. 43, 485–515 (2005).

17. Mirera, D. O. Status of the mud crab fishery in Kenya: A review. West. Indian Ocean J. Mar. Sci. 16, 35–45 (2017).

18. Zeng, Y. et al. Global potential and limits of mangrove blue carbon for climate change mitigation. Curr. Biol. 31, 1737–1743 (2021).

19. Menéndez, P., Losada, I. J., Torres-Ortega, S., Narayan, S. & Beck, M. W. The global flood protection benefits of mangroves. Sci. Rep. 10, 4404 (2020).

20. Rog, S. M., Clarke, R. H. & Cook, C. N. More than marine: revealing the critical importance of mangrove ecosystems for terrestrial vertebrates. Divers. Distrib. 23, 221–230 (2017).

21. Aburto-Oropeza, O. et al. Mangroves in the Gulf of California increase fishery yields. Proc. Natl. Acad. Sci. U. S. A. 105, 10456–10459 (2008).

22. Carrasquilla-Henao, M. & Juanes, F. Mangroves enhance local fisheries catches: A global meta-analysis. Fish Fish. 18, 79–93 (2017).

23. Tarimo, B., Winder, M., Mtolera, M. S. P., Muhando, C. A. & Gullström, M. Seasonal distribution of fish larvae in mangrove-seagrass seascapes of Zanzibar (Tanzania). Sci. Reports 2022 121 12, 1–13 (2022).

24. Dahdouh-Guebas, F. et al. Cross-cutting research themes for future mangrove forest research. Nat. Plants 8, 1131–1135 (2022).

25. Mukherjee, N. et al. The Delphi technique in ecology and biological conservation: Applications and guidelines. Methods Ecol. Evol. 6, 1097–1109 (2015).

26. Rovai, A. S. et al. Global controls on carbon storage in mangrove soils. Nat. Clim. Chang. 8, 534–538 (2018).

27. Twilley, R. R., Castañeda-Moya, E., Rivera-Monroy, V. H. & Rovai, A. Productivity and carbon dynamics in mangrove wetlands. in Mangrove Ecosystems: A Global Biogeographic Perspective: Structure, Function, and Services (eds. Rivera-Monroy, V., Lee, S., Kristensen, E. & Twilley, R.) 113–162 (Springer International Publishing, 2017).

28. Londoño, L. A. S., Leal-Flórez, J. & Blanco-Libreros, J. F. Linking mangroves and fish catch: A correlational study in the southern Caribbean Sea (Colombia). Bull. Mar. Sci. 96, 415–429 (2020).

29. Rodríguez-Rodríguez, J. A., Pineda, J. E. M., Trujillo, L. V. P., Rueda, M. & Ibarra-Gutiérrez, K. P. Ciénaga Grande de Santa Marta: The largest lagoon-delta ecosystem in the Colombian Caribbean. in The Wetland Book (eds. Finlayson, C., Milton, G., Prentice, R. & Davidson, N.) 1–16 (Springer, Dordrecht, 2016).

30. Freire, K. M. F. et al. Reconstruction of marine commercial landings for the Brazilian industrial and artisanal fisheries from 1950 to 2015. Front. Mar. Sci. 8, 659110 (2021).

31. Malik, A., Mertz, O. & Fensholt, R. Mangrove forest decline: Consequences for livelihoods and environment in South Sulawesi. Reg. Environ. Chang. 17, 157–169 (2017).

32. Silva, E. & Chiliquinga, J. S. Use of the mangrove red crab (Ucides occidentalis) in the Gulf of Guayaquil. Food Stud. 8, 53–69 (2018).

33. Nascimento, D. M. et al. Commercial relationships between intermediaries and harvesters of the mangrove crab Ucides cordatus (Linnaeus, 1763) in the Mamanguape River estuary, Brazil, and their socio-ecological implications. Ecol. Econ. 131, 44–51 (2017).

34. Pinheiro, M. A. A., Souza, M. R., Santos, L. C. M. & Fontes, R. F. C. Density, abundance and extractive potential of the mangrove crab, Ucides cordatus (Linnaeus, 1763) (Brachyura, Ocypodidae): subsidies for fishery management. An. Acad. Bras. Cienc. 90, 1381–1395 (2018).

35. Mota, T. A., Pinheiro, M. A. A., Evangelista-Barreto, N. S. & da Rocha, S. S. Density and extractive potential of “uçá”-crab, Ucides cordatus (Linnaeus, 1763), in mangroves of the “Todos os Santos” Bay, Bahia, Brazil. Fish. Res. 265, 106733 (2023).

36. Rönnbäck, P. The ecological basis for economic value of seafood production supported by mangrove ecosystems. Ecol. Econ. 29, 235–252 (1999).

37. Ahmad Adnan, N., Loneragan, N. R. & Connolly, R. M. Variability of, and the influence of environmental factors on, the recruitment of postlarval and juvenile Penaeus merguiensis in the Matang mangroves of Malaysia. Mar. Biol. 141, 241–251 (2002).

38. Flores, L., Licandeo, R., Cubillos, L. A. & Mora, E. Intra-specific variability in life-history traits of Anadara tuberculosa (Mollusca: Bivalvia) in the mangrove ecosystem of the Southern coast of Ecuador. Rev. Biol. Trop. 62, 473–482 (2014).

39. Bunting, P. et al. Global mangrove extent change 1996-2020: Global Mangrove Watch version 3.0. Remote Sens. 14, 3657 (2022).

40. Carrasquilla-Henao, M., González Ocampo, H. A., Luna González, A. & Rodríguez Quiroz, G. Mangrove forest and artisanal fishery in the southern part of the Gulf of California, Mexico. Ocean Coast. Manag. 83, 75–80 (2013).

41. Costa, R. C., Lopes, M., Castilho, A. L., Fransozo, A. & Simōes, S. M. Abundance and distribution of juvenile pink shrimps Farfantepenaeus spp. in a mangrove estuary and adjacent bay on the northern shore of São Paulo State, southeastern Brazil. Invertebr. Reprod. Dev. 52, 51–58 (2010).

42. Muyot, F. B., Magistrado, M. L., Muyot, M. C., Theresa, M. & Mutia, M. Growth performance of the mangrove red snapper (Lutjanus argentimaculatus) in freshwater pond comparing two stocking densities and three feed types. Philipp. J. Fish. 28, 1–17 (2020).

43. Malik, A., Fensholt, R. & Mertz, O. Economic valuation of mangroves for comparison with commercial aquaculture in south Sulawesi, Indonesia. Forests 6, 3028–3044 (2015).

44. Warningsih, T., Kusai, K., Bathara, L., Zulkarnain, Z. & Deviasari, D. Economic valuation of mangrove ecosystem services in Sungai Apit District, Siak Regency, Riau Province, Indonesia. IOP Conf. Ser. Earth Environ. Sci. 695, 012036 (2021).

45. Ickowitz, A., Lo, M. G. Y., Nurhasan, M., Maulana, A. M. & Brown, B. M. Quantifying the contribution of mangroves to local fish consumption in Indonesia: A cross-sectional spatial analysis. Lancet Planet. Heal. 7, e819–e830 (2023).

46. Santos, L. C. M., Pinheiro, M. A. A., Dahdouh-Guebas, F. & Bitencourt, M. D. Population status and fishery potential of the mangrove crab, Ucides cordatus (Linnaeus, 1763) in North-eastern Brazil. J. Mar. Biol. Assoc. United Kingdom 98, 299–309 (2018).

47. MacKenzie, C. L. The fisheries for mangrove cockles, Anadara spp, from México to Perú, with descriptions of their habitats and biology, the fishermen’s lives, and the effects of shrimp farming. Mar. Fish. Rev. 63, 1–39 (2001).

48. Gardner, C. J. et al. Value chain challenges in two community-managed fisheries in Western Madagascar: Insights for the small-scale fisheries guidelines. in The Small-Scale Fisheries Guidelines (eds. Jentoft, S., Chuenpagdee, R., Barragán-Paladines, M. J. & Franz, N.) 335–354 (Springer, 2017). doi:10.1007/978-3-319-55074-9_16.

49. Cannicci, S. et al. A functional analysis reveals extremely low redundancy in global mangrove invertebrate fauna. Proc. Natl. Acad. Sci. U. S. A. 118, e2016913118 (2021).

50. Hutchison, J., zu Ermgassen, P. S. E. & Spalding, M. D. The current state of knowledge on mangrove fishery values. in American Fisheries Society Symposium 83 3–15 (2015).

51. Adams, A. J. & Murchie, K. J. Recreational fisheries as conservation tools for mangrove habitats. in American Fisheries Society Symposium83 43–46 (2015).

52. Ndarathi, J., Munga, C., Hugé, J. & Dahdouh-Guebas, F. A socio-ecological system perspective on trade interactions within artisanal fisheries in coastal Kenya. West. Indian Ocean J. Mar. Sci. 19, 29–43 (2020).

53. FAO. ‘FAOSTAT Food Balances’ FAO. Retrieved from http://www.fao.org/faostat/en/#data/FBS. Accessed through Resource Watch, (16/06/2023). http://www.resourcewatch.org. (2020).

54. Robertson, W. D. & Piper, S. E. Population estimates of the crab Scylla serrata (Forskål, 1755) (Decapoda: Portunidae) in two closed estuaries in Natal, South Africa, from mark-recapture methods. South African J. Mar. Sci. 11, 193–202 (1991).

55. Likert, R. A technique for the measurement of attitudes. Arch. Psychol. 22, 55 (1932).

56. Stock, C. A. et al. Reconciling fisheries catch and ocean productivity. Proc. Natl. Acad. Sci. U. S. A. 114, E1441–E1449 (2017).

57. Polansky, L., Newman, K. B., Nobriga, M. L. & Mitchell, L. Spatiotemporal models of an estuarine fish species to identify patterns and factors impacting their distribution and abundance. Estuaries and Coasts 41, 572–581 (2018).

58. R Core Team. R: A Language and Environment for Statistical Computing. (2023).

59. Bivand, R. S., Pebesma, E. J. & Gomez-Rubio, V. Applied Spatial Data Analysis with R, Second Edition. (Springer, 2013).

60. Hijmans, R. J. raster: Geographic Data Analysis and Modeling. R package version 3.6-20. https://CRAN.R-project.org/package=raster. (2023).

61. Pierce, D. ncdf4: Interface to Unidata netCDF (Version 4 or Earlier) Format Data Files. R package version 1.17. https://CRAN.R-project.org/package=ncdf4. (2019).

62. Bivand, R. S., Keitt, T. & Rowlingson, B. gdal: Bindings for the ‘Geospatial’ Data Abstraction Library. R package version 1. 5-23. (2021).

63. Bradley, M., Nagelkerken, I., Baker, R. & Sheaves, M. Context dependence: A conceptual approach for understanding the habitat relationships of coastal marine fauna. Bioscience 70, 986–1004 (2020).

64. Primavera, J. H. Mangroves as nurseries: Shrimp populations in mangrove and non-mangrove habitats. Estuar. Coast. Shelf Sci. 46, 457–464 (1998).

65. Worthington, T. A. et al. A global biophysical typology of mangroves and its relevance for ecosystem structure and deforestation. Sci. Rep. 10, 14652 (2020).

66. Saintilan, N. Relationships between estuarine geomorphology, wetland extent and fish landings in New South Wales estuaries. Estuar. Coast. Shelf Sci. 61, 591–601 (2004).

67. Manson, F. J., Loneragan, N. R., Harch, B. D., Skilleter, G. A. & Williams, L. A broad-scale analysis of links between coastal fisheries production and mangrove extent: A case-study for northeastern Australia. Fish. Res. 74, 69–85 (2005).

68. Henderson, C. J., Gilby, B. L., Stone, E., Borland, H. P. & Olds, A. D. Seascape heterogeneity modifies estuarine fish assemblages in mangrove forests. ICES J. Mar. Sci. 78, 1108–1116 (2021).

69. Reis-Filho, J. A., Giarrizzo, T. & Barros, F. Tidal migration and cross-habitat movements of fish assemblage within a mangrove ecotone. Mar. Biol. 163, 1–13 (2016).

70. Sheaves, M., Johnston, R. & Baker, R. Use of mangroves by fish: New insights from in-forest videos. Mar. Ecol. Prog. Ser. 549, 167–182 (2016).

71. Lee, S. Y. Relationship between mangrove abundance and tropical prawn production: A re-evaluation. Mar. Biol. 145, 943–949 (2004).

72. Carrère, L., Lyard, F., Cancet, M. & Guillot, M. FES 2014, a new tidal model on the global ocean with enhanced accuracy in shallow seas and in the Arctic region. in EGU General Assembly 2015, held 12-17 April, 2015 in Vienna, Austria (2015).

73. Reis-Filho, J. A., Harvey, E. S. & Giarrizzo, T. Impacts of small-scale fisheries on mangrove fish assemblages. ICES J. Mar. Sci. 76, 153–164 (2019).

74. Stewart, K. R. et al. Characterizing fishing effort and spatial extent of coastal fisheries. PLoS One 5, e14451 (2010).

75. Saintilan, N., Hossain, K. & Mazumder, D. Linkages between seagrass, mangrove and saltmarsh as fish habitat in the Botany Bay estuary, New South Wales. Wetl. Ecol. Manag. 15, 277–286 (2007).

76. Goodridge Gaines, L. A. et al. Linking ecosystem condition and landscape context in the conservation of ecosystem multifunctionality. Biol. Conserv. 243, 108479 (2020).

77. Olson, J. C. et al. Recruitment of juvenile snapper (Lutjanidae) in the Middle Florida Keys: Temporal trends and fine-scale habitat associations. Gulf Caribb. Res. 35, GCFI1–GCFI13 (2024).

78. Baker, R., Sheaves, M. & Johnston, R. Geographic variation in mangrove flooding and accessibility for fishes and nektonic crustaceans. Hydrobiologia 762, 1–14 (2015).

79. Daqamseh, S. T., Al-Fugara, A., Pradhan, B., Al-Oraiqat, A. & Habib, M. MODIS derived sea surface salinity, temperature, and chlorophyll-a data for potential fish zone mapping: West Red Sea coastal areas, Saudi Arabia. Sensors 19, 2069 (2019).

80. Fernandes, J. A. et al. Projecting marine fish production and catch potential in Bangladesh in the 21st century under long-term environmental change and management scenarios. ICES J. Mar. Sci. 73, 1357–1369 (2016).

81. Boutin, J. et al. ESA Sea Surface Salinity Climate Change Initiative (Sea_Surface_Salinity_cci): Monthly sea surface salinity product, v03.21, for 2010 to 2020. (2021).

82. Watson, R. & Pauly, D. Systematic distortions in world fisheries catch trends. Nature 414, 534–536 (2001).

83. Putri, R. et al. The relationship between small pelagic fish catches with sea surface temperature and chlorophyll in Makassar Strait waters. J. Iktiologi Indones. 22, 65–76 (2022).

84. Merchant, C. J. et al. Satellite-based time-series of sea-surface temperature since 1981 for climate applications. Sci. Data 6, 1–18 (2019).

85. Ningsih, W. A. L., Lestariningsih, W. A., Heltria, S. & Khaldun, M. H. I. Analysis of the relationship between chlorophyll-a and sea surface temperature on marine capture fisheries production in Indonesia: 2018. IOP Conf. Ser. Earth Environ. Sci. 944, 012057 (2021).

86. Behrenfeld, M. J. & Falkowski, P. G. Photosynthetic rates derived from satellite-based chlorophyll concentration. Limnol. Oceanogr. 42, 1–20 (1997).

87. Worthington, T. A. et al. A global biophysical typology of mangroves version 3. (2023) doi:10.5281/zenodo.8340259.

88. Stammer, D. et al. Accuracy assessment of global barotropic ocean tide models. Rev. Geophys. 52, 243–282 (2014).

89. Ray, R. D., Eanes, R. J., Egbert, G. D. & Pavlis, N. K. Error spectrum for the global M2 ocean tide. Geophys. Res. Lett. 28, 21–24 (2001).

90. Pinheiro, J., Bates, D., DebRoy, S., Sarkar, S. & R Core Team. nlme: Linear and Nonlinear Mixed Effects Models. R package version 3. 1-152, https://CRAN.R-project.org/package=nlme. https://cran.r-project.org/package=nlme (2021).

91. Zuur, A. F., Ieno, E. N., Walker, N., Saveliev, A. A. & Smith, G. M. Mixed Effects Models and Extensions in Ecology with R. (Springer, 2009).

92. Kaschner, K. et al. AquaMaps: Predicted range maps for aquatic species. World wide web electronic publication, https://www.aquamaps.org. Version 08/2015. https://www.aquamaps.org (2015).

93. Ragionieri, L., Fratini, S. & Schubart, C. D. Revision of the Neosarmatium meinerti species complex (Decapoda: Brachyura: Sesarmidae), with descriptions of three pseudocryptic Indo-West Pacific species. Raffles Bull. Zool. 60, 71–87 (2012).

94. Prener, C., Grossenbacher, T. & Zehr, A. biscale: Tools and Palettes for Bivariate Thematic Mapping. R package version 1.0.0, https://CRAN.R-project.org/package=biscale. (2022).

