## Supporting Information for "The global fish and invertebrate abundance value of mangroves"

### Supplemental information

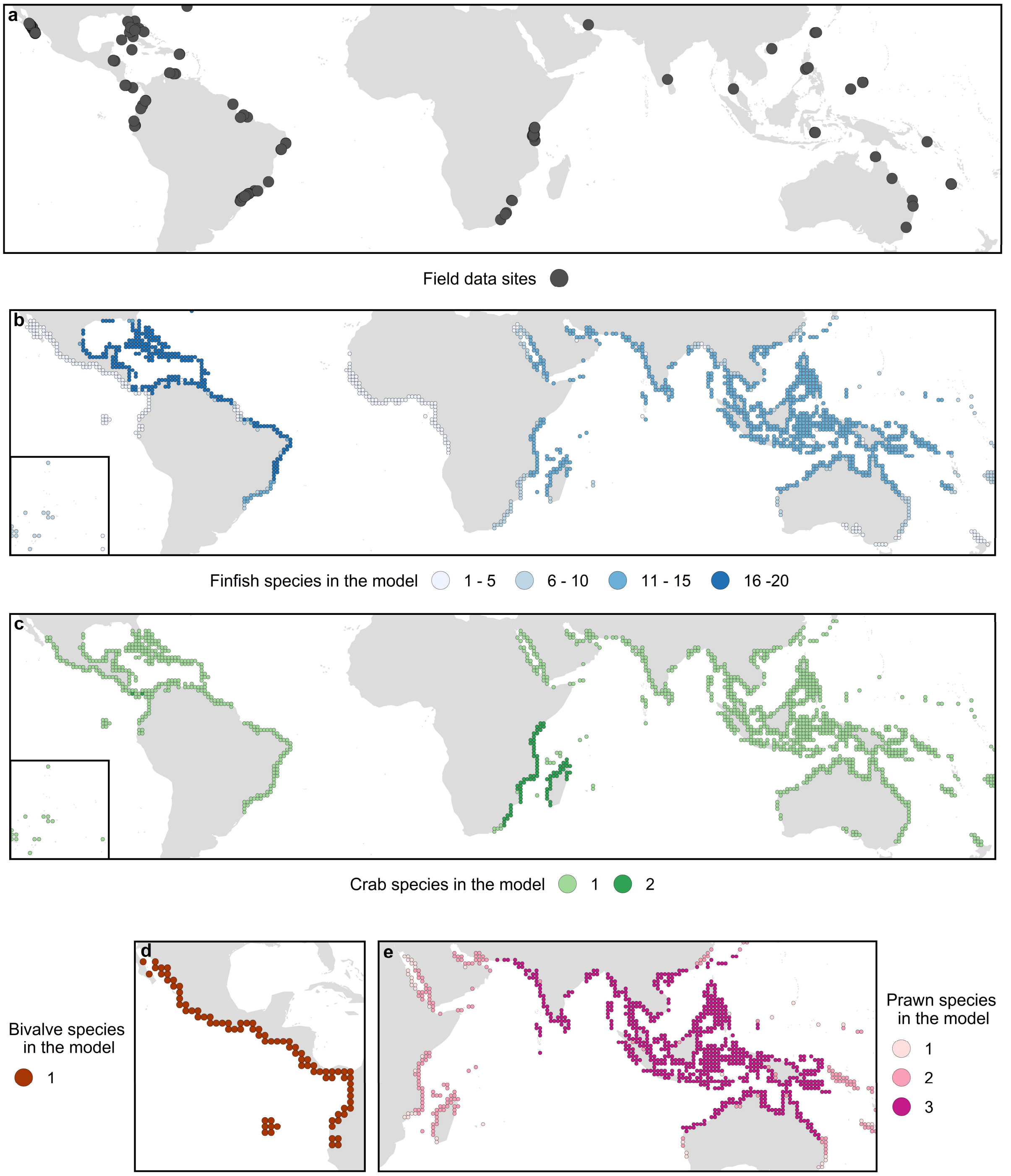


**Fig. S1.** a) Location of the 481 field measurements of fish and invertebrate density, and the number of b) finfish c) bivalve, d) prawn and e) crab species included in the model. Data summarised within 1° cells, inset shows Hawaii and Pacific islands.

**Table S1.** The species included in the model.

| Species | Group |
|---|---|
| *Anadara tuberculosa* | Bivalve |
| *Neosarmatium africanum* | Crabs |
| *Scylla serrata* | Crabs |
| *Ucides cordatus* | Crabs |
| *Ucides occidentalis* | Crabs |
| *Atherinomorus lacunosus* | Finfish |
| *Caranx latus* | Finfish |
| *Centropomus undecimalis* | Finfish |
| *Cetengraulis edentulus* | Finfish |
| *Chaetodipterus faber* | Finfish |
| *Chanos chanos* | Finfish |
| *Diapterus auratus* | Finfish |
| *Gerres cinereus* | Finfish |
| *Gerres filamentosus* | Finfish |
| *Haemulon flavolineatum* | Finfish |
| *Haemulon sciurus* | Finfish |
| *Lutjanus analis* | Finfish |
| *Lutjanus apodus* | Finfish |
| *Lutjanus argentimaculatus* | Finfish |
| *Lutjanus argentiventris* | Finfish |
| *Lutjanus cyanopterus* | Finfish |
| *Lutjanus fulviflamma* | Finfish |
| *Lutjanus griseus* | Finfish |
| *Lutjanus jocu* | Finfish |
| *Lutjanus russellii* | Finfish |
| *Lutjanus synagris* | Finfish |
| *Monodactylus argenteus* | Finfish |
| *Mulloidichthys martinicus* | Finfish |
| *Scarus guacamaia* | Finfish |
| *Siganus canaliculatus* | Finfish |
| *Sillago sihama* | Finfish |
| *Sparisoma rubripinne* | Finfish |
| *Sphyraena barracuda* | Finfish |
| *Terapon jarbua* | Finfish |
| *Penaeus indicus* | Prawns |
| *Penaeus merguiensis* | Prawns |
| *Penaeus monodon* | Prawns |

**Table S2.** Factors considered by experts during the Delphi process. Experts were asked to identify which factors they considered to be important in explaining the density of fish and invertebrates in mangroves.

| **Variable** |
|---|
| Annual mean air temp (°C) |
| Annual maximum air temp (°C) |
| Annual minimum air temp (°C) |
| Annual mean sea surface temp (°C) |
| Air temp annual range (°C) |
| Annual maximum sea surface temp (°C) |
| Annual minimum sea surface temp (°C) |
| Sea surface temp annual range (°C) |
| Seasonality (contrast between seasons) |
| Mangrove extent (area in ha) |
| Mangrove edge (length in metres) |
| Mangrove above ground biomass (tonnes per ha) |
| Mangrove species richness |
| Mangrove typology/ecomorphology (e.g., riverine, fringe, lagoonal) |
| Extent of estuary/embayment associated with mangroves (area in ha) |
| Latitude |
| Longitude |
| Freshwater volume entering the mangrove (mean annual freshwater flows) |
| Salinity (ppt) |
| Duration of inundation/flooding of mangrove area (months per year) |
| Duration of inundation/flooding of mangrove area (hours per day) |
| Tidal regime (micro (<2m), meso (2-4m), macro (>4m) tides)) |
| Coastal bathymetry |
| Total suspended particulate matter (coastal turbidity from satellite imagery; $g/m^3$) |
| Coastal primary productivity (standing stock Chla from satellite imagery; $mg/m^3$) |
| Oceanic primary productivity (standing stock Chla from satellite imagery; $mg/m^3$) |
| Nitrate concentration (micromol/l) |
| Phosphate concentration (micromol/l) |
| Underlying geology of the system (e.g., carbonate versus acidic rocks) |
| Fish and invertebrate inter-species differences in dependency on mangroves (an index of the relative importance of mangrove for each species' life history) |
| Fish and invertebrate species probability of occurrence (species specific maps of probability of occurrence) |
| Degree of artisanal fishing (national weighting of prevalence of artisanal fishing) |
| Size of coastal pop. (no. of people within a given distance of mangrove) |
| Size of rural coastal population (non-urban population within a given distance of mangrove) |
| Access to market (distance to market and density of infrastructure) |
| Mangrove condition (degree of forest loss 2000-2012) |
| Mangrove protected area designation (areas within protected areas versus areas with no protection) |
| Biogeographic region (to capture variation not explained by other variables listed over and above the differences which result from values represented in other variables) |
| Proximity to other habitats (seagrass) |
| Proximity to other habitats (coral reef) |
| Proximity to other habitats (shallow unstructured habitat i.e. mud or sand) |
| Dissolved Oxygen |

| |
|---|
| Fishing pressure |
| Land use adjacent to mangrove |
| Water quality |
